# Neuraminidase Inhibition Contributes to Influenza A Virus Neutralization by Anti-Hemagglutinin Stem Antibodies

**DOI:** 10.1101/397976

**Authors:** Ivan Kosik, Davide Angeletti, James S. Gibbs, Matthew Angel, Kazuyo Takeda, Martina Kosikova, Vinod Nair, Heather D. Hickman, Hang Xie, Christopher C. Brooke, Jonathan W. Yewdell

## Abstract

Broadly neutralizing antibodies (Abs) that bind the influenza virus hemagglutinin (HA) stem may enable universal influenza vaccination. Here, we show that anti-stem Abs sterically inhibit viral neuraminidase activity against large substrates, with activity inversely proportional to the length of the fibrous NA stalk that supports the enzymatic domain. By modulating NA stalk length in recombinant IAVs, we show that anti-stem Abs inhibit virus release from infected cells by blocking NA, accounting for their *in vitro* neutralization activity. NA inhibition contributes to anti-stem Ab protection in influenza infected mice, likely due at least in part to NA-mediated inhibition of Fc*γ*R dependent activation of innate immune cells by antibody bound to virions. FDA approved NA inhibitors enhance anti-stem based Fc*γ* dependent immune cell activation, raising the possibility of therapeutic synergy between NA inhibitors and anti-stem mAb treatment in humans.

## Introduction

The severe 2017-18 influenza season provides a timely if unwelcome reminder of the limitations of current vaccines in coping with antigenic drift in the viral surface proteins, hemagglutinin (HA) and neuraminidase (NA). HA mediates viral attachment to cell surface sialic acid residues and subsequent fusion of viral and cellular membranes. NA releases nascent virus from infected cells by removing terminal sialic residues from glycoproteins and glycolipids.

Current vaccines induce antibodies (Abs) specific for the HA head. Head-binding antibodies neutralize IAV infectivity *in vitro* by blocking virus attachment, and depending on the epitope recognized, by preventing the conformational alterations needed to trigger membrane fusion. Due to the immunodominance of the HA head following infection and standard vaccination, Ab responses drive rapid evolution in the head that enables viral escape. In contrast to the HA head, the stem domain is highly conserved and cross-reactive between strains within the same group. Early proof-of principle studies find that a stem-specific mAb could block HA mediated fusion, neutralize IAV and protect against IAV disease in animal models (Okuno et al., 1993; Okuno et al., 1994; Russ et al., 1987; Styk et al., 1979). Further studies from many labs indicate that broadly neutralizing (BN) stem-specific Abs are common in humans and can be induced by standard vaccination protocols, albeit at low levels relative to head-specific Abs (Chiu et al., 2015; Corti et al., 2017; Henry Dunand and Wilson, 2015). Much more robust BN stem-specific responses can be elicited by native stem only immunogens (Impagliazzo et al., 2015; Lu et al., 2014; Mallajosyula et al., 2014; Yassine et al., 2015) or a prime-boost head-stem chimeric HA molecule strategy (Krammer et al., 2017).

Although Ab-driven antigenic drift in the stem may ultimately limit stem-based vaccination (Doud et al., 2018; Lees et al., 2014), as a promising “universal” vaccine strategy it is critical to understand the mechanism(s) of anti-stem Abs in reducing/preventing IAV disease in humans. Stem binding Ab have been reported to block viral entry into cells by preventing the acid-induced conformations alterations in HA required to catalyze viral-cell membrane fusion (Vareckova et al., 2003), and to prevent release from infected cells, through an unknown mechanism (Yamayoshi et al., 2017). *In vivo*, anti-stem Abs may largely exert protection via Fc mediated activation of humoral and cellular innate immune functions (Cox et al., 2016; DiLillo et al., 2014a)

HA is typically present on virions at more than 5-fold higher molar amounts than NA. Abs specific for the HA globular as well as for stem domain can sterically interfere with NA activity against large protein substrates as long as virus remains intact (Kosik and Yewdell, 2017; Rajendran et al., 2017; Russ et al., 1974), suggesting a possible mechanism for stem-Ab mediated inhibition of viral release from infected cells. Here we show that NA inhibition can be a major contributor of BN activity of stem-specific Abs *in vitro* and *in vivo*, in the latter case likely by interfering with Fc-receptor activated innate immune cell anti-viral effector activity.

## Results

### BN HA stem specific mAbs inhibit viral neuraminidase

The commonly used ELLA method for measuring Ab-mediated NA inhibition (NAI) is complicated by blockade of viral access to the plate-bound substrate by anti-HA head Abs that block viral attachment (Kosik and Yewdell, 2017). This does not, however, apply to stem-binding Abs which do not block virus attachment even at saturating concentrations (Figure S1A). Anti-stem mAbs 310-16G8 and 310-18F8 (Whittle et al., 2014) efficiently inhibit N1 or N2 NAs using viruses with H1 HA stems (PR/8/34 (H1N1), Cal/4/09 (H1N1), chimeric cH5/1N2 (H5headH1 stem N2)) (Fig. 1). Similarly, we found that the CR8020 mAb that binds H3 HA (group 2) stems could block N2 NA in several H3N2 strains, and for FI6, also N1 in PR8, demonstrating that NAI can be mediated by a group I/II cross-reactive stem-specific Ab (Fig. 1).

**Figure 1.**
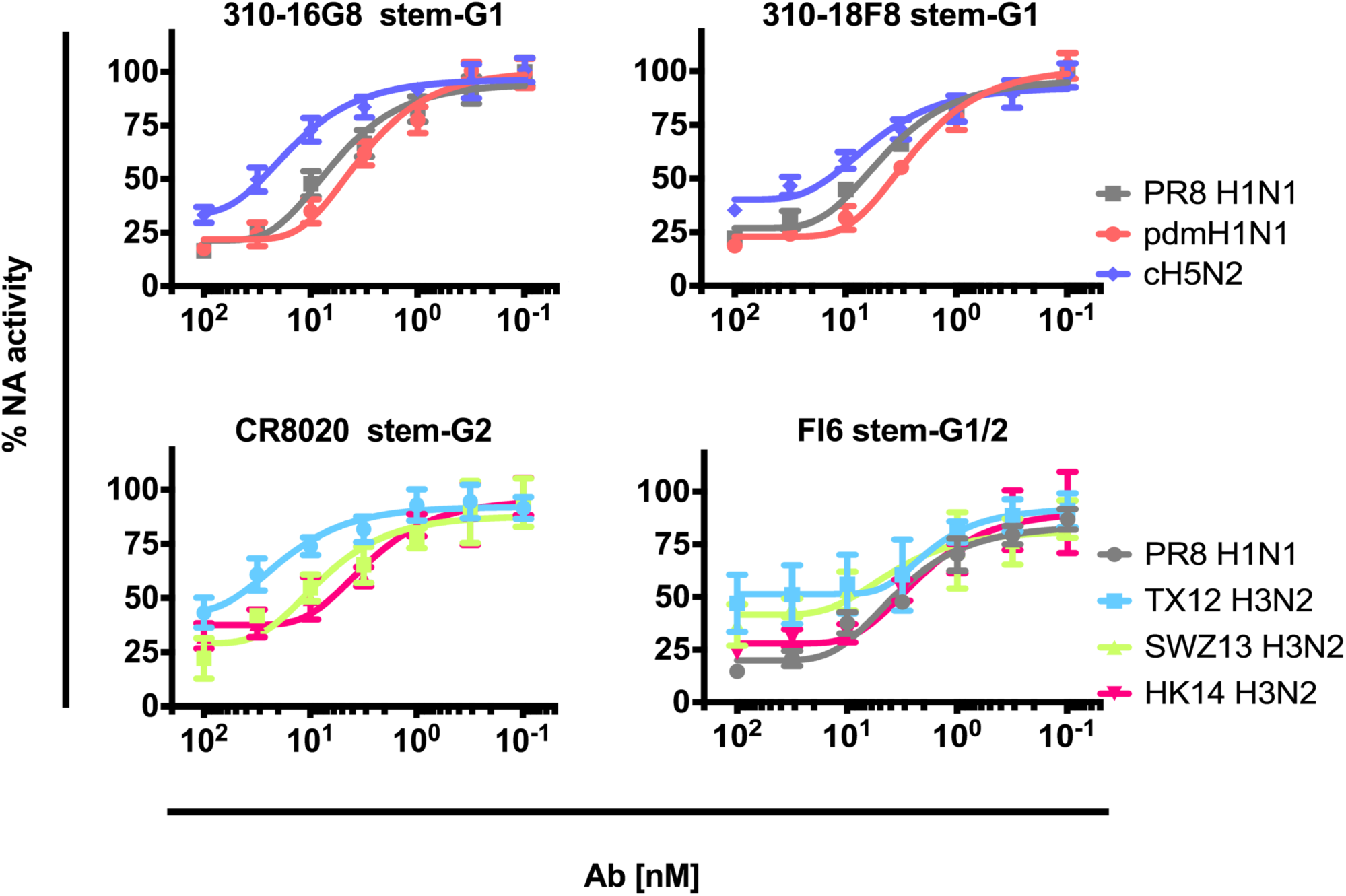
BN anti-HA stem mAbs sterically block NA activity. We measured the capacity of purified anti-stem mAbs on NA activity via ELLA against the viruses indicated. ELLA is based on NA removal of terminal sialic acids from plate bound fetuin, which is assessed by binding of a lectin for the exposed penultimate galactose. mAbs used are specific for either group 1 (G1), group 2 (G2) HA stems, or bind both groups (G1/2). We normalized data by NA activity in the absence of the Ab (set to 100%). Error bars indicate the SD of duplicate samples acquired in three independent experiments.

As with globular domain specific mAbs (Kosik and Yewdell, 2017), NAI with stem binding Abs was abrogated using detergent-treated virus, illustrating the steric nature of inhibition (Fig. S1B).

### Neuraminidase inhibition of stem specific Abs is inversely related to NA stalk length

Curiously, the chimeric H5/1N2 virus exhibits 2-3 fold increased resistance to stem mAb-mediated NAI compared to PR8, despite sharing the identical stem sequence (Fig.1). Sequence alignment of the relevant N1 and N2 NAs revealed a 15-amino acid extension in the N2 NA stalk region (Fig. S1C), suggesting that NA stalk length may influence the NAI activity of anti-stem Abs. The NA stalk exhibits natural length variability (Li et al., 2011) and its modulation is frequently associated with host adaptation and pathogenicity changes (Bi et al., 2015; Matsuoka et al., 2009; Sun et al., 2013).

To examine the influence of NA stalk length on anti-stem Ab function, we generated recombinant PR8 viruses with NAs lacking a stalk (del24) or with a 20-residue insertion (ins20) (Fig. S2A). Despite no obvious differences in the morphology of purified virions by cryoEm (Fig. S2B), immunoblotting for NA confirmed the expected alteration in M_r_ in SDS-PAGE (Fig. S2C). Using two independently generated recombinant virus preps we found that wt, del24 and ins20 purified virions contained similar amounts of NA and HA by quantitative immunoblotting (Fig. S2C). Kinetic analysis of NA activity in purified viruses using the small fluorogenic substrate 4-methylumbelliferyl-a-D-N-acetylneuraminic acid (MUNANA) (Table S1) revealed that adding or deleting stalk amino acids had minor effects on NA V_max_ or Km. By contrast, lengthening the stem greatly accelerated NA-mediated release from RBCs while shortening the stem retarded release (Fig. S2D), confirming previous findings (Chockalingam et al., 2012). While ins20 and del24 exhibit altered release from RBCs compared to wt, all the viruses replicated with similar kinetics in MDCK SIAT1 cells, consistent with similar replication also *in vivo* (Fig. S2D). We next examined HA stem-Ab mediated NAI using the stalk mutants.

Using ELISA, we established that NA stalk length does not affect the binding of anti-HA stem mAbs 310-16G8 and 310-18F8 to plate-bound purified virus (Fig. S3 upper panels). Similarly, flow cytometry revealed that the NA stalk does not affect stem Ab binding to HA on infected MDCK cells (Fig. S3). Despite this, shortening the NA stalk increased the NAI activity of the same mAbs by ∼4-fold relative to wt virus while lengthening the stalk had the oppositeeffect, resulting in a 10-20-fold difference in NAI titers in ins20 vs. del24 viruses (Fig. 2A).

**Figure 2.**
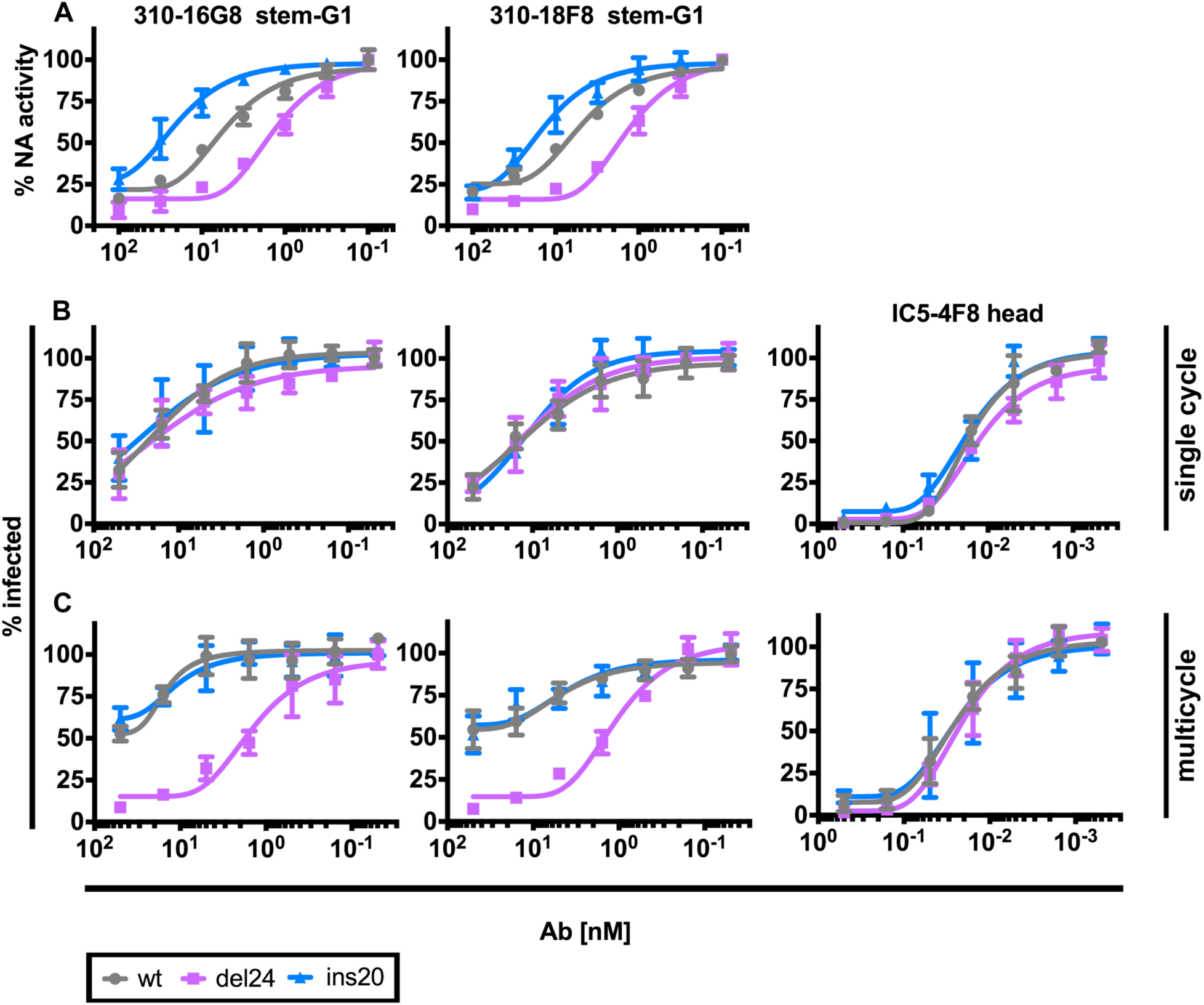
Stalk length governs anti-stem Ab NAI and multicycle VN activity. A. NAI activity of the indicated purified mAbs determined by ELLA as in Figure 1 using PR8 and mutants engineered to add 20 residues to the NA stalk (ins20) or deleted the stalk (del24).
B. VN activity of the indicated purified mAbs determined by incubating mAbs with the mCherry reporter viruses indicated prior to adding to MDCK SIAT1 cells and after 4.5 hours, quantitating infected cells by flow cytometry to detect mCherry expression. Data are normalized with 100% set as the fraction of infected cells in the absence of Ab.
C. As in B but measuring infected cells 18h post infection to allow multicycle infection. Error bars indicate the SD of duplicate samples acquired in three independent experiments.

### NA inhibition contributes to virus-neutralizing activity of stem-specific Abs by inhibiting release

Our findings predict that stem-binding Abs should neutralize viral infectivity by inhibiting NA-mediated release from infected cells. To test this prediction, we examined the effect of NA stalk length on mAb neutralization. For this, we generated a panel of reporter viruses that express mCherry, enabling precise flow cytometric enumeration of virus-infected cells. We added virus preincubated with mAbs at 37°C to MDCK SIAT1 cells and measured frequency of IAV infected cells 4 h post infection (p.i.). IC5-4F8, a control mAb specific for the Sb antigenic site in the head domainof H1 neutralized viruses equally regardless of NA stalk (Fig. 2B). Neutralization by two stem-specific Abs was much less efficient (even when factoring in Ab avidity) but did not vary with NA stalk length after single replication cycle (Fig. 2B).

By contrast, when infection time was extended by 14 h to enable multicycle infection both stem specific mAbs demonstrated a ∼ 100-fold increase in neutralization against del24 relative to wt or ins20 (Fig. 2C). This effect was stem-specific, as it was not observed with IC5-4F8 (Fig. 2C).

A simple explanation for the NA-specific effect by anti-stem Abs in multistep neutralization of del24 is that anti-stem Abs block virus release, retarding cell to cell transmission. We therefore added mAbs 4 h after infecting MDCK-SIAT1 cells with wt, del24 or ins20 PR8, and measured NA activity of virus released into the media from cells over the next 6 h using NA activity with MUNANA as a proxy. We confirmed that mAbs did not alter the infection *per se* by quantitating intracellular NP and NA in fixed and permeabilized cells by indirect immunoassay (Fig. S4). Both stem mAbs tested blocked release of each virus (Fig. 3). By contrast, the HA head-specific mAb-IC5-4F8 as expected (or a control mAb specific for an irrelevant Ag) did not block virus release (Fig. 3, Fig. S4). Importantly, both stem mAbs tested exhibited greatly enhanced effects on del24 release from cells (Fig. 3C), providing an explanation for the enhanced activity of anti-HA stem Abs in the multicycle neutralization assay (Fig. 2C).

**Figure 3.**
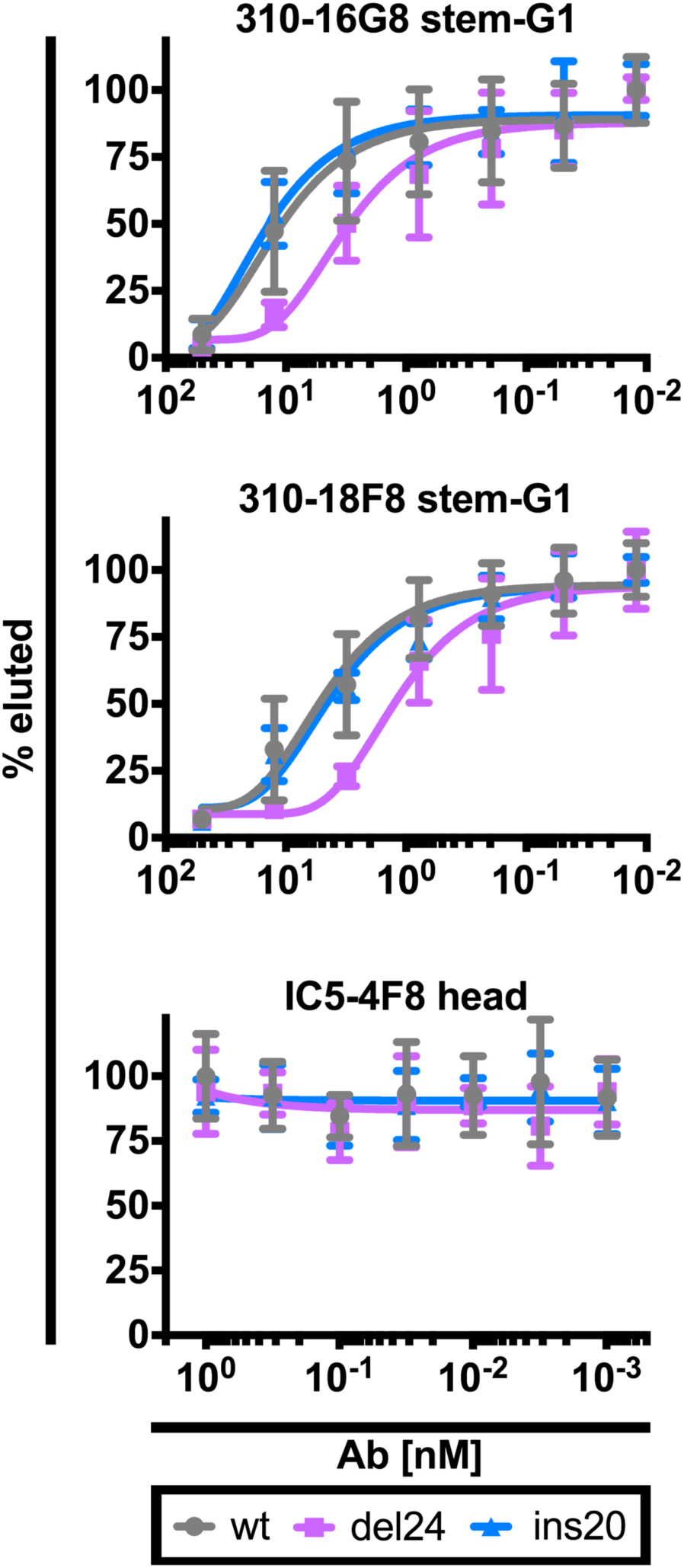
Anti-stem Abs inhibit nascent virus release from infected cells. Four hr p.i. of MDCK SIAT1 cells with the viruses indicated we added graded amounts of the indicated mAbs. Six hr later, we collected the supernatant and used NA activity against a fluorescent substrate to quantitate released virions. The fluorescent signal in absence of Ab was set as 100% for each virus. Error bars indicate the SD of duplicate samples acquired in up to four independent experiments.

### NA stalk length controls effectiveness of HA stem specific Abs in limiting IAV pathogenesis

Extending these findings to an *in vivo* model of IAV pathogenesis, we passively transferred stem or head specific mAbs into C57Bl/6 mice, 4 h later challenged with pathogenically equal high doses of wt or NA stalk variant viruses and monitored weight loss. Following infection without transferred Ab, 100% of mice were humanely sacrificed between d 4 and 7 p.i. when they reached >30% weight loss. As expected, mice receiving the HA head-specific mAb IC5-4F8 were efficiently protected against each virus, with minimal weight loss and 100% survival (Fig. 4A). Surprisingly, in mice passively administered 310-16G8, infection with either wt or del24 resulted in delayed and attenuated weight loss and all mice recovered from by 10 d p.i. (Fig 4A). In contrast, mice infected with ins20 receiving 310-16G8 lost weight with similar kinetics as control, PBS administered mice, with only 17% of ins20-infected mice recovering (Fig. 4A). Higher doses of wt virus resulted in more rapid weight loss and 100% mortality in 310-168-treated mice infected with wt virus, but 80% protection and recovery by day 13 in del24-infected Ab-treated mice (Fig. 4B).

**Figure 4.**
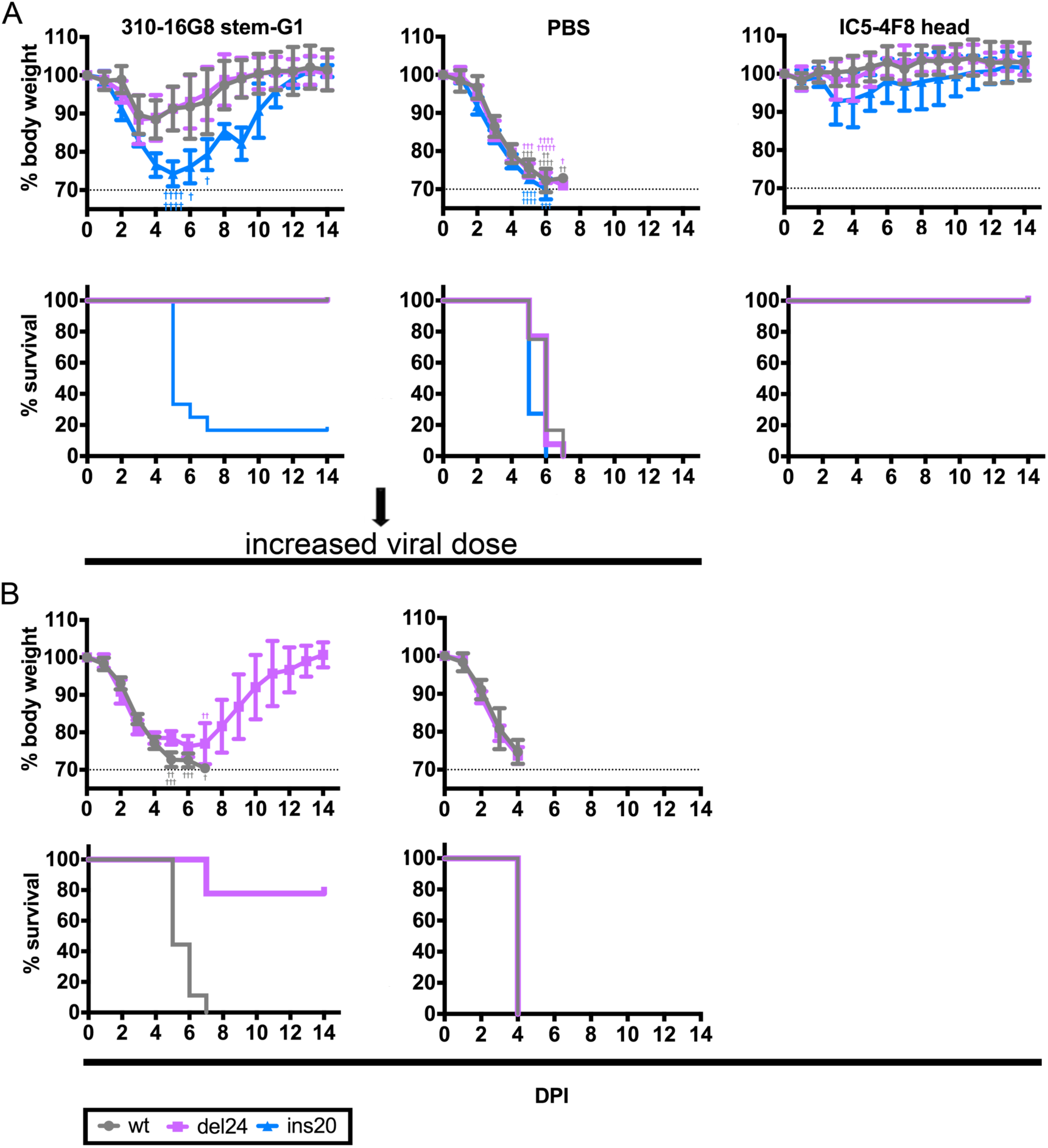
Protection mediated by anti-stem Abs is dependent on NA inhibition. A. We passively transferred mice (n=10-13 mice/group, 5mg/kg) with the stem specific mAb 310-16G8. PBS (negative control) or head specific mAb IC5-4F8 (positive control). Four h post transfer we infected mice intranasally with pathologically equal doses of wt, del24 and ins20 viruses. We monitored morbidity (body weight) and mortality (human sacrifice at >30% weight loss) for fourteen days.
B. As in panel A, but with 9 mice per group infected with 4-fold more wt or de24 viruses. As a control we passively transferred mice (n=4/group) with PBS only. Error bars indicate the standard deviation (SD). Presented results are sum of three independent experiments (n=3-4/group)

Histological staining of lung sections on d4 p.i. revealed less tissue destruction and immune infiltration in IC5-4F8 *vs.* PBS treated animals infected with each of the viruses, as expected (Fig. 5A). There were only minor differences between wt and ins20-infected mice in terms of perivascular inflammation and bronchial epithelial damage as well as in alveolar damage and immune cell infiltration in 310-16G8 treated mice between wt and ins20 viruses (Fig. 5A left/right field), despite the clearly increased morbidity of ins20 infection in these mice. The only statistically significant difference in histopathological scoring was a decreased immune cell infiltration in alveoli of del24-infected mice (Fig. 5B), clearly shown in the insets. Fluorescent immunostaining of the IAV NP in 310-16G8 treated mice mirrors alveolar histopathological burden with decreased frequency of NP positive cells in particular for del24 infected mice (Fig. 5C,D). It is clear, that NA stem length, and therefore NA activity against sterically-susceptible substrates, can greatly affect the *in vivo* activity of anti-HA stem Abs, decreasing alveolar damage in parallel with reducing virus infected alveolar cells.

**Figure 5.**
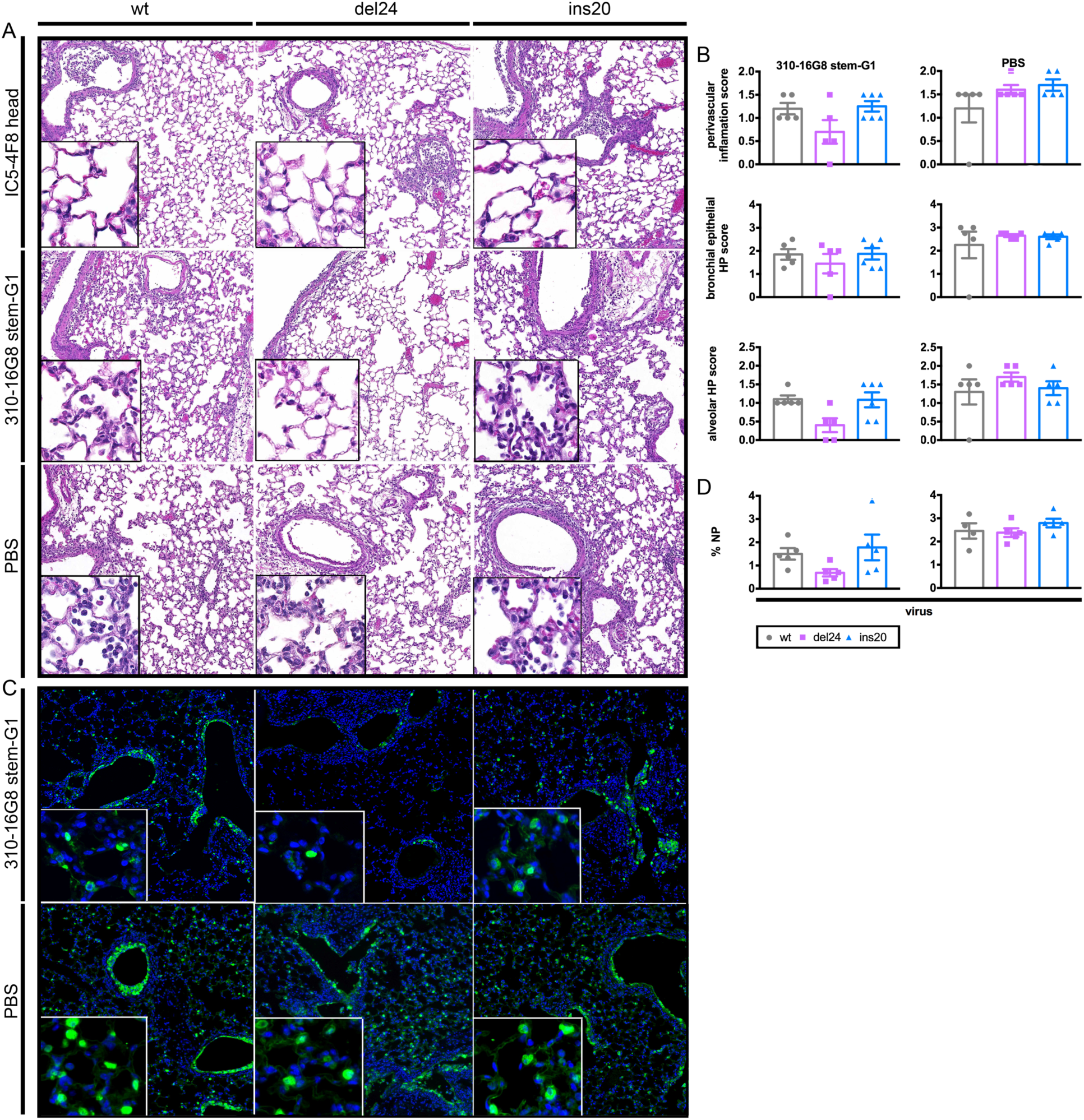
Pathogenesis modultated by anti-stem Abs is dependent on NA inhibition. We passively transferred mice (n=5-6/group, 5mg.kg^−1^) with stem-specific mAb 310-16G8 or head-specific mAb IC5-4F8, or PBS. Four h post transfer we infected mice intranasally with pathologically equal doses of wt, del24 and ins20 viruses, four d post challenge, we collected lungs, and stained sections with hematoxylin (dark purple-nucleoli) and eosin (pink/red-cytoplasm) to visualize lung tissue damage and immune cell infiltration.

A. Representative lung section fields of mice infected with indicated viruses and treated as indicated
B. Quantitation of histopathogical changes
C. Representative fields showing influenza NP expression revealed by immunofluorescence.
D. Quantitation of NP staining using Bitplane Imaris 9.1 software to determine the fraction of infected cells, using DNA staining to identify all cellular nuclei as a denominator.

Error bars indicate the SD of duplicates-triplicates samples acquired in two independent experiments.

### NA Inhibits Fc receptor based innate cell activation

Despite the clear virus neutralizing activity of anti-HA stem Abs *in vitro*, several studies indicate that their protective effect in mice depends on their interaction with Fc *γ* receptors (Fc*γ*R), implicating the required participation of Fc*γ*R bearing innate immune cells (NK cells, macrophages, dendritic cells and others) or/and complement. HA-stem specific Abs are known to activate cells via Fc*γ*R interaction in a process that requires HA binding to SA residues on the Fc*γ*R expressing cell (DiLillo et al., 2016; Dilillo et al., 2014b), since it is blocked by head-specific mAbs that block virus attachment.

We could confirm these findings using Jurkat reporter cells engineered to express luciferase after engagement of Ab-Ag complexes with the Fc*γ*RIIIa receptor expressed from a transgene (Jurkat cells are human T cell leukemia that do not naturally express Fc receptors or other NK activating receptors). Reporter cells were activated in a viral infection and stem Ab dependent manner. The addition of IC5-4F8, which binds non-competitively with stem Abs in close proximity to the HA receptor binding site, completely blocked activation (Fig. 6A).

**Figure 6.**
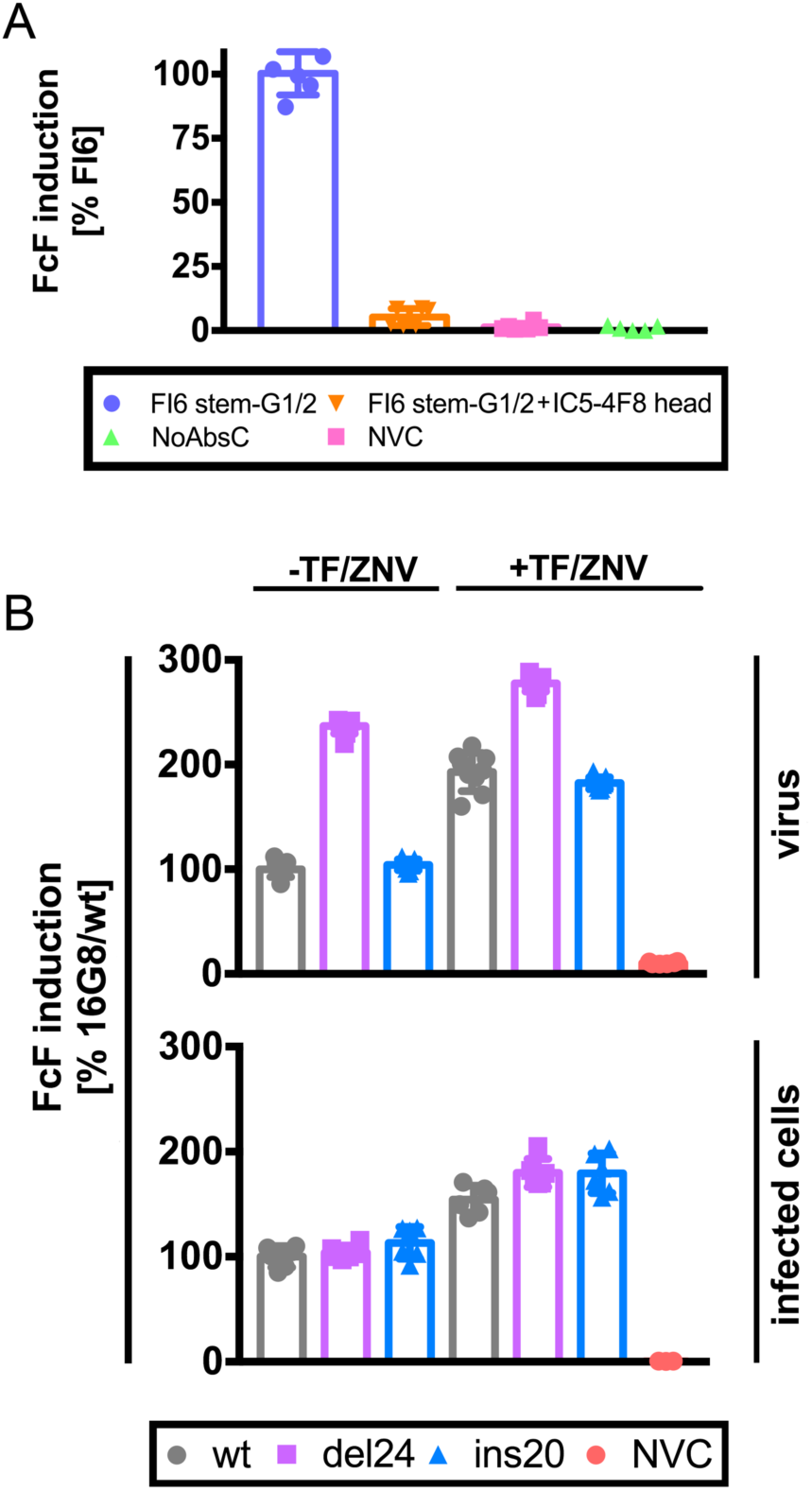
NA inhibition governs anti-stem HA Fc mediated cellular activation by virus, but not infected cells. A. We infected MDCK SIAT1 cells with PR8 and at 5.5 hr p.i. added the mAbs indicated and Jurkat Fc*γ*R-reporter cells. After overnight incubation, we measured luciferase expression to gauge cell activation. 100% activation is defined by FI6 mAb alone. Error bars indicate the SD of triplicates samples acquired in two independent experiments.
B. In the top graph, we pre-incubated wt, del24 or ins20 viruses (normalized by HAU) with the 16G8 anti-stem mAb and added Jurkat reporter cells in the presence or absence of oseltamivir (tamiflu) and zanamivir (TF/ZNV). In the bottom graph, we infected cells at equal m.o.i. with the viruses indicated and 5-6 h p.i. added 16G8 and Jurkat reporter cells in the presence and absence of the TF/ZNV. After overnight incubation, we measured luciferase expression to gauge cell activation. 100% activation is defined by FI6 mAb alone. Error bars indicate the SD of triplicate samples acquired in two independent experiments.

Having established a robust system for measuring stem-Ab based Fc*γ*R coupled innate cell activation we next examined whether the anti-NA activity of stem-specific Abs enhances activation. After incubating reporter cells overnight with virus-Ab mixtures, cellular activation was enhanced implicating NA-based 2-fold by del24 relative to wt or ins20 virus, inhibition of Fc*γ*R-based activation. Consistent with this observation, adding FDA approved NA inhibitors (oseltamivir and zanamivir) increased reporter cell activation in all circumstances (Fig. 6B top panel). The greatest enhancement was observed with wt and ins20 virusesin the presence of NA inhibitors (Fig. 6B). Curiously, despite what should be complete NA inhibition with the drugs, activation remained less than observed with del24, or del24 + NA inhibitors, which achieved maximal activation. NA inhibitors also enhanced anti HA-stem Ab-dependent innate cell activation by MDCK SIAT1 cells infected with wt or and NA stalk variant viruses. Interestingly however, NA stalk length did not modulate activation (Fig. 6B bottom panel).

Our interpretation of these data are that first, virus or cell surface NA interferes with Fc*γ*R-based cell activation by cleaving SA on reporter cells, which can be blocked with NA inhibitors. Secondly, stem Ab-based steric effects on NA inhibition are far less sensitive on infected cells *vs.* virus, probably due to the relaxed organization of HA and NA spikes on the cell surface relative to the highly organized geometry on virions.

## Discussion

HA stem-specific Abs are perhaps the most promising approach for improving the duration and effectiveness of influenza vaccination. It is therefore important to better understand how anti-stem Abs provide *in vivo* protection. Evidence from mouse passive Ab transfer studies indicates a critical role for Fc*γ*Rs in protection at lower (i.e. more physiological) stem Ab concentrations, implicating innate cell mechanisms. These could include NK cell mediated killing of infected cells (Bar-On et al., 2013), macrophage clearance of virus, and cytokine/chemokine secretion by any of the many type of Fc*γ*R expressing cells.

Here we demonstrate a potentially critical role for NA in the anti-viral activity of HA specific Abs. By modulating the anti-NA effect of stem Ab using NA stalk deleted and extended viruses, we could show that stem binding Abs inhibit NA activity against large substrates, thereby blocking nascent virus release from infected cells. This phenomenon was reported previously (Yamayoshi et al., 2017), however the mechanism has not been defined.

Our findings have interesting implications for the organization of glycoproteins on the virion surface. We unequivocally demonstrate that anti-stem Ab have sufficient access to HA on intact virions to inhibit NA activity, extending cryoelectron microscopy-based observations that Abs can access HA stem on intact virions (Harris et al., 2013; Wasilewski et al., 2012; Williams et al., 2017). Examination of the Ab NAI titration curves (Fig. 2) reveals several intriguing features.

First, in agreement with Rajendran et al., at anti-stem Ab saturation ∼ 25% of NA activity remains. This suggests that this fraction of NA is sufficiently physically segregated from HA on virions to prevent steric inhibition from neighboring HA. Decreasing stalk length reduces the resistant fraction by ∼50%. As NA is generally present in clusters on the virion surface (Calder et al., 2010; Harris et al., 2006; Wasilewski et al., 2012), the resistant fraction may represent NA on the interior of clusters of 4 or more molecules. If shortening stalk length decreased cluster size, this could account for the decreased resistant fraction.

Second, as NA stalk length increases, the amount of anti-stem Ab needed to block NA activity increases up to 20-fold, despite stalk length having no apparent effect on anti-HA binding to virions. As the Ab concentration increases, monovalent interactions will be favored, perhaps enabling more Abs bound per NA cluster, thus increasing the reach of stem-bound Abs.

In as much as the *in vivo* activity of stem-binding Abs is based on Fc*γ*R ligation (DiLillo et al., 2016; Dilillo et al., 2014b; Mullarkey et al., 2016), such activities would not be expected to contribute to Ab based protection *in vivo*. Consistent with this conclusion, we only detected up to three-fold decrease in viral replication in stem Ab treated mice, despite a clear protective effect. While *in vivo* viral replication was shown correlate with stem-Ab dose, protection was also reported at low doses. In such a case viral replication can be reduced less than 10-fold (Dilillo et al., 2014b; Kallewaard et al., 2016) while still completely protecting against mortality.

Remarkably, extending the NA stalk greatly reduced the protective effect of anti-HA stem mAbs, strongly implicating NA inhibition in Ab-mediated protection in our experimental system (Fig. 4, 5). Conversely, shortening the NA stalk reduced immune alveolar inflammation and lymphocyte infiltration, and increased protection at high virus doses. These effects are likely due to modulating NA activity. Indeed, we show that NA inhibits Fc*γ*R reporter cell activation as clearly revealed by the enhancing effect of small molecule NA inhibitors.

NA inhibition of Fc*γ*R expressing cells is consistent with the finding that anti-HA head-specific Abs block NK cell ADCC killing while NA-specific Abs enhance it (He et al., 2016). The simplest explanation for this phenomenon is that HA-mediated binding to SA on innate immune cells is required for cell activation. Indeed, Ab-independent NK cell recognition of IAV-infected cells requires HA binding to NK activating receptors NKp46, NKp44 (Mandelboim et al., 2001), 2B4 or NTB (Duev-Cohen et al., 2016), and viral NA inhibits this activation (Bar-On et al., 2013). Because the Jurkat Fc*γ*R reporter cells we and others (He et al., 2016) have used are not known to express NK activating receptors, it appears that their requirement for HA binding in Ab-based activation reflects HA binding to other cell surface ligands.

NA blockade of Ab-dependent and independent innate immune cell anti IAV activity supports the possibility that NA inhibitors owe their clinical efficacy to more than just blocking virus release, as suggested by Bar-On *et al.* (Bar-On et al., 2013). Further, if innate immune effector functions limit IAV transmission, NA-mediated antagonism could contribute to NA evolution, with NA stalk length possibly being under selective pressure by virtue of its functional interaction with HA-stem specific Abs. Pragmatically, the ability of NA inhibitors to enhance Fc*γ*R-mediated innate immune cell activation of stem-Abs bound to viruses or infected cells suggests the possible clinical synergism between NA inhibitors and anti-stem mAbs in humans.

## Material and methods

### Animals

We purchased C57BL/6 mice from Taconic Farm. We used female 8-to 12-week-old mice randomly assigned to experimental groups. We held mice under specific pathogen-free conditions. All animal procedures were approved and performed in accordance with the NIAID Animal Care and Use Committee Guidelines. For passive transfer, we weighed and injected mice with 5mg/kg of each mAb in 400 μl of sterile saline solution intraperitoneally. For mouse infections, we anesthetized mice with isoflurane (3.5%) and injected 25 μl of virus intranasally (wt=3.5×10^3^, del24=5×10^3^, ins20=4×10^3^) diluted in sterile saline supplemented with 0.1% bovine serum albumin (BSS-BSA). We recorded weight for 14 days and euthanized mice when mice reached 30% weight loss. We excluded mice (n=2) that showed no signs of infection and weight loss compared to rest of the animals in given experiment. When needed, we harvested and homogenized lungs and determined viral titers.

### Cell lines and viruses

We cultured Madin-Darby Canine Kidney cells SIAT1(MDCK SIAT1) in Dulbecco’s Modified Eagle Medium (DMEM, Gibco) supplemented with 8% FBS (HyClone) and 500 μg/ml gentamicin (Quality Biologicals) and 500 μg/ml geneticin (Gibco) at 37°C and 5% CO2. We cultured human embryonal kidney 293T cells(HEK293T) in Dulbecco’s Modified Eagle Medium (DMEM, Gibco) supplemented with 8% FBS (HyClone) and 500 μg/ml gentamicin at 37°C and 5% CO2. We used Expi293 cells (Thermo Fisher) for BN HA stem mAbs expression. We propagated and cultured the cells as recommended by the manufacturer. We employed the following viruses in this study. The A/Puerto Rico/8/34/H1N1(Walter Gerhart), A/California/07/2009/H1N1, A/Texas/50/2012/H3N2, A/Switzerland/9715293/2013/H3N2, A/Hong Kong/4801/2014/H3N2, A/Puerto Rico/8/34/H1N1/ins20 NA, A/Puerto Rico/8/34/H1N1/del24 NA, chimeric HA influenza virus with HA head domain from the A/Vietnam/1203/04 (H5N1), HA stem domain from PR8 (Hai et al., 2012) and NA from A/Udorn/72/H3N2 (cH5/1N2). We generated the NA stalk viruses by co-transfection of the eight influenza gene coding plasmids (generously provided by Dr. Adolfo Garcia-Sastre; Icahn School of Medicine at Mt. Sinai, New York) as published previously (Kosik et al., 2018). To generate deletion in the NA stalk while preserving cysteine at position 49 we performed mutagenic overlapping PCR with forward primer: 5’AGCCATTCA TGC AAGGACACAACTTCAGTGATATTAACC’3 and reverse primer: 5’TGTGTCCTTGCATGAATGGCTAATCCATATT GAG’3. We used the pDZ-NA plasmid as template along with Kod Hot start DNA polymerase (Millipore Sigma). For the NA insertion variant, we used 5 prime phosphorylated forward primer: 5’AATTTTCTTACTGAAAAAGTTGTTGCTGGGAAGGACACAACTTCAGTGATATTAACCG’3 containing 30 nucleotides from NA stalk region of the A/Goose/Guangdong/1/96/H5N1 (AF144304): and reverse primer: 5’GGTATTGCTGATGTTGACATATGTTTGATTTACCCAGGTGCTATTTTTATAGGTAATG’3 containing 15 nucleotides from NA stalk region of the A/Goose/Guangdong/1/96/H5N1 and 15 nucleotides from A/WSN/33/H1N1 (LC333187). We followed manufacturer recommendations to PCR mutagenize the pDZ-NA ORF. We recircularized linearized plasmid PCR products with Rapid DNA Ligation Kit (Roche), transformed plasmids into DH5*α* competent cells (Thermo Fisher). After purification with plasmid purification (Qiagen), we sequenced plasmids. To rescue chimeric cH5/1N2, we co-transfected plasmid harboring chimeric HA with N2 plasmid (generously provided by Dr. Peter Palese and Dr. Kanta Subbarao respectively) combined with PR8 core plasmids and followed as described above. For the self-reporting NS1-mCherry expressing virus we first modified reverse genetics system pDZ plasmid as follows.

We cloned NS1-mCherry into the pDZ vector plasmid in several steps. First, we amplified NS1 by PCR with primers NS-5’-SapI (GCTCTTCAGGGAGCAAAAGCAGGGTGACAAAG) and NS1-mCherry bot (GCCGCTGCCATCGATGCCAACTTCTGACCTAATTGTTC). We PCR amplified mCherry with primers NS1-mCherry top (TGGCATCGATGGCAGCGGCATGGTGAGCAAGGGCGAGGAG)and mChe-2A-NEP bot (AGGCTAAAGTTGGTCGCGCCACCGCTGCCCT TGTACAGCTCGTCCATGCC). We PCR amplified NEP with primers NEP-exon-1 (GAAAACCCGGGCCCGATGGATCCAAACACTGTGTCAAGCTTTCAGGACATACTGCTGAGGATG TCAA), which encodes NEP exon 1 fused to exon 2, and primer NS-3’-SapI(GCTCTTCTATTAGTAGAAACAAGGGTGT TTT). After all amplification steps, we dissected PCR fragments from an agarose gel and purified with the QIAquick gel extraction kit (Qiagen). We extended NEP fragment further using primers 2A-NEP (GGCGCGACCAACTTTAGCCTACTGAAACAGG CGGGCGATGTGGAAGAAAACCCGGGCCCGATGGA) and NS-3’-SapI, which added the “self-cleaving” 2A sequence of porcine teschovirus. We diluted purified mCherry and 2A-NEP fragments 1:50 and joined with splice-overlap extension PCR using primers NS1-mCherry top and NS-3’-SapI. We digested the NS1 fragment and mCherry-2A-NEP fragment with SapI and ClaI restriction enzymes and ligated to pDZ vector digested with SapI. The final construct codes for NS1 fused to a six amino acid linker (GIDGSG) fused to mCherry followed by the 2A sequence and both exons of NEP. We sequenced the final plasmid and used it instead of pDZ-NS in rescue as described above. We propagated viruses in ten days old embryonated chicken eggs for 48 hours and we stored allantoic fluid at −80°C. For virus purification, we clarified allantoic fluid at 4700 RPM for 10 min, and pelleted virus by centrifugation for 2 hr at 27,000 RPM. We incubated pellets overnight in 2 ml PBS with calcium and magnesium (PBS++) and purified virus by centrifugation on a discontinuous 15–60% sucrose gradient, collecting virus at the interface, and pelleting 34,000 RPM for 2 hr After resuspending the pellet in 500 µl PBS++ overnight we measured total viral protein with the DC Protein Assay (Bio-Rad).

### Viral sequencing

We extracted influenza virus genomic RNA from clarified allantoic fluid using the QIAamp Viral RNA Mini Kit (Qiagen) according to the manufacturer’s protocol. We amplified complete genomes with a multiplex RT-PCR protocol using SuperScript III One-Step RT-PCR System with Platinum Taq DNA Polymerase, and primers targeting conserved 5’ and 3’ sequences in the viral RNA (Zhou and Wentworth, 2012). Libraries for next generation sequencing were constructed from 1ng amplified genomes using Nextera XT DNA Library Preparation Kit (Illumina), and sequenced using the MiSeq Reagent Kit v2 on the MiSeq Platform (Illumina). Viral sequences were assembled de novo using a high-throughput assembly pipeline (Mena et al., 2016), and variant statistics were assembled using custom perl scripts.

### Antibodies

We produced the human IgG1 bnHA stem mAbs 310-16G8, 310-18F8, FI6, CR8020 (Ekiert et al., 2011; Whittle et al., 2014) by transient plasmid (kindly provided by Dr. Adrian McDermott) co-transfection in the Expi293 Expression System (Thermo Fisher) following manufacturer recommendations. The IC5-4F8 Sb specific, H2-6A1 Sa specific, HA2 specific mAb RA5-22, NP specific mAb HB65, rabbit anti C-terminus NP polyclonal serum 2364 (487-498aa), NA specific NA2-1C1 mAb, rabbit anti C-terminus NA polyclonal serum, TW1.3 vaccinia specific mAb were prepared in the lab as published previously (Kosik and Yewdell, 2017; Yewdell et al., 1981). We prepared the polyclonal anti-HK68 mouse serum by single dose intramuscular immunization (5 μg) of whole purified UV-inactivated virus. We collected serum at 21 d post immunization. After heat inactivation at 56°C for 30 min we stored serum at −80°C.

### Virus hemagglutination titer

We diluted allantoic fluid containing half log serial dilutions of virus in DPBS in round bottom ninety-six-well plate (Greiner Bio-One). We combined 50 μl of the virus dilutions with 50 μl 0.5% turkey red blood cells and incubated at 4°C for one hour. We determined HA titer as reciprocal of highest dilution providing full hemagglutination. We tested all viruses in triplicates in two independent experiments.

### In Cell Immunoassay and TCID_50_

We seeded MDCK SIAT1 cells (10,000-50,000 per well) in ninety-six-well plate (Costar). The following day we washed cells with DPBS twice and added virus diluted in infection media (MEM medium (Gibco) containing 0.3% BSA (fraction V, Roche), 10 mM HEPES (Corning), 500 μg/ml gentamicin (Quality Biologicals) and 1 μg/ml TPCK trypsin (Worthington)). At times p.i. indicated we fix-permeabilized the cells with 100% ice cold methanol for 20 min. We washed the cells with DPBS and we added 30 μl of HB65 anti NP mAb or/and NA2-1C1 anti NA mAb (1 μg/ml) diluted in Odyssey^®^ Blocking Buffer PBS (OBB) (Li-Cor) for one hour at room temperature. We washed the plate with DPBS containing 0.05% NP-40 (Thermo Fisher) twice and we added IRDye^®^ 800CW goat anti mouse or IRDye^®^ 680RD Goat anti-Rabbit secondary antibody (Li-Cor) 1000x diluted in OBB. After one-hour incubation at room temperature we washed the plate twice as described above with one final milliQ water washing step. For TCID_50_ determination, we prepared half log_10_ dilutions of the virus sample in infection media. We transferred virus dilutions in replicates of eight to MDCK SIAT1 cells in ninety-six-well plate (Costar). At 18hr p.i. we proceed as described above. We used the HB65 anti NP specific mAb along with 1,000fold diluted IRDye^®^ 800CW goat anti mouse secondary antibodies to detect IAV positive wells. We scanned the plates by near-infra red imaging system OdysseyCLx (Li-Cor). We used ImageStudioLite software to quantify the measured signal. For the positive/negative threshold we used doubled averaged signal of eight uninfected wells. We used the Spearman and Karber method to calculate TCID_50_. For quantification of NA and NP expression we subtracted the average signal of four uninfected wells. We plotted TCID_50_, NA and NP signal data using GraphPad Prism7.

### Protein gels and immunoblotting

We diluted freshly purified virus samples (25μg/ml) in DPBS and combined three volumes of virus with one volume of NuPAGE™ LDS Sample Buffer (4X) (Thermo Fisher) containing 100mM DTT. We heated samples at 70°C for 10 min, cooled them on ice and loaded 5 μl (100ng) of the sample on NuPAGE 4-12% Bis-Tris Protein Gel (Thermo Fisher), separating proteins by electrophoresis with Chameleon Pre-Stained Protein Ladder (Li-Cor) on 4–12% Bis-Tris Gels (Invitrogen) at 150 V for 90 min. We transferred proteins to nitrocellulose membranes using an iBlot device at P3 setting for 7 min. We blocked membranes with OBB for one hour at room temperature and incubated with rabbit anti C-terminus NA polyclonal serum (5,000x dilution) and RA5-22 mAb (1 μg/ml) diluted in OBB. We washed the membrane in DPBS containing 0.05% NP-40 three times and incubated for one hour at room temperature with IRDye^®^ 800CW goat anti mouse or IRDye^®^ 680RD Goat anti-Rabbit secondary antibody (Li-Cor) diluted 20,000-fold in OBB. We washed the membrane three times with one additional milliQ water washing step. We scanned the membrane with the near-infra red imaging system OdysseyCLx (Li-Cor). We used ImageStudioLite software to quantify signal and plotted the data using GraphPad Prism7 software.

### ELISA

We coated ELISA plates (half area ninety-six-well, Greiner Bio-One) with 10ng of respective purified IAV diluted in DPBS (50 μl per well). After overnight incubation at 4°C, we washed plates three times with DPBS supplemented with 0.05% Tween-20. We serially diluted (half log dilutions) Abs starting from 10-100 nM in 1% BSA in DPBS, and added 50 μl to plates for 90 min at 37°C. After extensive washing with DPBS+0.05% Tween-20, we detected bound Abs by incubating plates with 50 μl HRP-conjugated rat anti-mouse IgG kappa chain (Southern Biotech) for 1 h at room temperature. We washed ELISA plates with DPBS+0.05% Tween-20 and incubated with SureBlue TMB Microwell Peroxidase Substrate (KPL) for 5 min at room temperature. We stopped the enzymatic reaction by adding HCl. We determined the A_450_ on a Synergy H1 plate reader (Biotek) and calculated the Kd from dilution curves using GraphPad Prism 6 software to fit one site binding.

### ELLA

We performed ELLA as described previously (Kosik and Yewdell, 2017). We avoided Tween-20 in all steps to preserve virion integrity. We determined the virus dilution yielding an A_450_ between 1 and 1.5. We also normalized samples to the a similar HAU titer. We coated half area ninety-six-well ELISA plates (Greiner Bio-One) with 50 μl fetuin solution in DPBS (25 μg/ml) overnight at 4°C. We diluted Abs initially to 200 nM and then serially by 3.1-fold. We pre-incubated 25 μl of Ab dilutions with 25 μl virus in sample diluent (DPBS, 1%BSA) for 60 min at 37°C and then added the samples to fetuin coated plate for 18-22 h at 37°C. We washed plates extensively with washing buffer (DPBS, 0.05% Tween-20) and then added 50 μl peanut-HRP (Sigma Aldrich) diluted 1,000-fold in sample diluent for one hour at room temperature in the dark. After washing we detected and read the plate as for ELISA. To liberate HA and NA from virion we added 0.5% TritonX-100 in sample diluent for 10 min at room temperature, and then followed the standard ELLA procedure. We normalized data by signal in absence of mAb. We plotted data with GraphPad Prism7 software and fit nonlinear regression curves using the dose response inhibition model.

### Attachment inhibition (AI) assay

We performed AI assay as described previously (Kosik and Yewdell, 2017). Briefly, we coated black, half area high binding 96-well plates (Nunc) with 50 μl fetuin in DPBS (25 μg/ml) overnight at 4°C. We determined the highest dilution of the virus yielding sufficient signal. We diluted mAbs or H3N2 specific mouse sera initially to 200 nM or 20-fold respectively and then serially by half log_10_ steps. We mixed Abs with diluted allantoic fluid and incubated for 60 min at 37°C. We then transferred 50 μl of Ab-virus mixture to washed fetuin-coated plates, incubated for 1h at 4°C, and removed free virus by washing 6-times with DPBS. We added 50 μl of 200 μM MUNANA in 33mM MES pH 6.5 with 4mM CaCl_2_ and incubated for 1 hour at 37°C. We measured fluorescence (Ex=360nm, Em=450nm) using a Synergy H1 plate reader (Biotek). The signal in the absence of Ab defines 100% attachment. We fitted nonlinear regression curves using the dose response inhibition model with GraphPad Prism7 software.

### Flow cytometry

For flow cytometry-based neutralization assay we seeded 50,000 MDCK SIAT1 cells to ninety-six-well tissue culture treated plates (Costar). On the next day, we pre-incubated at 37°C for one hour 50 μl of half log_10_ serially diluted Abs with 50 μl of the respective mCherry expressing virus at MOI=0.05-0.1 per well. We diluted Abs and virus in infection media. We transferred virus-Ab mixture to washed MDCK SIAT1 cells and incubated samples at 37°C for the time indicated (4-5 h/18 h). We washed cells twice with DPBS and added 30 μl of 0.005% trypsin-EDTA (Gibco) for 20 min, 30 μl BSS-BSA and then 30 μl 1.6% paraformaldehyde (PFA) (Electron Microscopy Science) for 20 min. We sedimented cells by centrifugation at 1,500 RPM for 5 min and washed samples with BSS-BSA two times. We also included uninfected and samples without Ab as controls. For comparing the access of the HA stem epitope in the context of cell expressed HA, we seeded 2.5 million MDCK SIAT1 cells per T25 tissue culture flasks and on the next day infected cells with viruses (moi=1) diluted in infection media. 18 hr p.i., we washed cells twice with DPBS, detached from the flask with trypsin-EDTA (Gibco) and combined with half log_10_ serial dilutions of mAbs. We incubated samples at 37°C for one hour, washed with BSS-BSA and combined with 50 μl 200x diluted mouse anti-human FITC (Pharmigen) or 5,000x diluted donkey anti-mouse AF488 (Jackson Immunology). We washed samples three times, fixed with 4% PFA for 20 min washed three times again and resuspended in BSS-BSA. Samples were analyzed using a BD LSRFortessa X-20 instrument. Analysis was performed using FlowJo software (TreeStar). We normalized the frequencies of mCherry positive cells by the non-antibody treated samples and fitted nonlinear regression curves using the dose response inhibition model for neutralization assay or one site binding for epitope access assay with GraphPad Prism7 software.

### Cell surface virus elution assay

We seeded 20,000 MDCK SIAT1 cells to ninety-six-well tissue culture treated plates (Costar). The next day, we washed cells twice with DPBS and infected cells with viruses (moi=5) diluted in infection media without trypsin. At 4hr p.i. we washed cells with DPBS twice and added 50 μl of serially diluted (4fold) mAbs. At 10hr p.i. we transferred 40 μl of the supernatant to round bottom ninety-six-well plate (Costar) and pelleted the cells at 4,700 RPM for 10 min. We gently transferred 30 μl of the supernatant to half area black ninety-six-well plate (Costar) and combined with 30 μl of 200 μM MUNANA in 33mM MES pH 6.5 with 4mM CaCl^2^. We incubated the samples for 1 hour at 37°C. We measured fluorescence (Ex=360nm, Em=450nm) using a Synergy H1 plate reader (Biotek). The signal in the absence of Ab defines 100% elution. We fitted nonlinear regression curves using the dose response inhibition model with GraphPad Prism7 software.

### ADCC assay

We seeded 10,000 MDCK SIAT1 cells to ninety-six-well white tissue culture treated plates (Costar). The following day we washed cells twice with DPBS and infected cells with viruses (moi=5) as described above. At 6 hr p.i. we combined 25 μl of 200nM mAb with 25 μl of ADCC-RL cells (Jurkat cell line expressing luciferase gene under control of the NFAT response element and stably expressing human Fc*γ*RIIIa V158) (Promega) containing 50,000 in RPMI 1640 medium with 4% low IgG serum (Promega). We added the Ab-ADCC-LR mixture onto the MDCK SIAT1cells (E:T=5:1). After overnight incubation at 37°C in 5% CO_2_, we added 50μl of Bright-Glo™ Luciferase Assay lysis/substrate buffer (Promega) and measured luminescence within 5 min using a Synergy H1 plate reader (Biotek). Alternatively, we combined 25 μl of respective virus containing 40 HAU with 25 μl of 300 nM mAb, incubated virus-Ab mixtures at 37°C for one hour and added 25 μl ADCC-LR (50,000). We incubated mixtures overnight at 37°C in 5% CO_2_ and measured luminescence signal as described above. We included samples in the absence of the virus as negative controls. When indicated, in addition to mAbs, mixture of the oseltamivir phospahe (TF, American Radiolabeled Chemicals) and zanamivir hydrate (ZNV, Moravek inc) were added to samples at 2.5 μM final concentration. We normalized the data to the signal of FI6 or 310-16G8 on wt PR8 virus. We plotted the data using GraphPad Prism7 software.

### NA catalytic activity

We diluted freshly purified viruses (∼2 μg/ml) in 33mM MES pH 6.5 with 4mM CaCl_2_ to contain equal NA amount based on multiple immunoblot experiments. We twofold-serially diluted MUNANA in the same buffer starting at 500 μM and pre-incubated virus solutions and MUNANA dilutions at 37°C for 30 min to equilibrate temperatures. We combined 20 μl of virus with 20 μl of respective MUNANA dilution in half area ninety-six-well plate (Costar) and incubated the samples at 37°C to generate fluorescent product of the catalytic reaction. After 60 min incubation we measured fluorescence generated (Ex=360nm, Em=450nm) using a Synergy H1 plate reader (Biotek). We measured background fluorescence of respective MUNANA dilutions and subtracted it from samples. In parallel, we diluted 4-MU (the catalytic product of MUNANA cleavage) serially (twofold) and we measured fluorescence as described above. We generated calibration curve by fitting linear regression in GraphPad Prism7 software and we interpolated amount of 4-MU produced by viruses. We fit the data to Michaelis-Menten regression in GraphPad Prism7 software.

### RBCs virus release assay

We measured relative NA activity of the viruses in allantoic fluid. We normalized samples based on equal NA activity (determined by MUNANA) and we diluted samples serially (twofold) in PBS++ and combined 50 μl of the virus with 50 μl of 0.5% turkey red blood cells diluted in PBS++. We incubated plates at 4°C for one hour, recorded the HA titer and transferred the plates to 37°C. We recorded complete elution every 30 min for seven hours. We divided complete elution dilution at time indicated by HA titer to normalize for viral particles amount. We transformed the data as % of wt PR8 elution at 7 hours. We plotted the data in GraphPad Prism7 software.

### 2D Cryo electron microscopy

For CEM, we froze 3.5μl of viral suspension on a glow discharged 200 mesh R2/2 Quantifoil™ Cu grids (Quantifoil, Großlöbichau, Germany) using Leica EM GP plunge freezer (Leica Microsystems Inc., Wetzlar, Germany). We blotted the grids for 2s at 99% relative humidity using 1s wait and drain time respectively. We mounted plunge frozen grids in autogrids assembly and imaged under cryo conditions at 300kV on Titan Krios (ThermoFisher Scientific, Hillsboro, OR) using Falcon II direct electron detector. We collected Cryo 2D Images using a volta phase plate for increased contrast at a nominal magnification of 29K which corresponds to a pixel size of 2.87Å with 1 micron defocus at a total dose of 12 e^−^/Å^2^.

### Histopathology and Immunohistochemistry

We infected mice intranasally with PR8 influenza A wt, del24 or ins20 NA variant viruses (3,000-5,000 TCID_50_). At 4 days p.i., we sacrificed the mice and inflated lung by 10% buffered formalin through the trachea We fixed lungs overnight and processed for paraffin embedding. We made serial sections at 5 um thickness and stained every 10th slides by Hematoxylin and Eosin. We dewaxed the unstained and stained with influenza NP antibody (rabbit, 172) followed with Alexa 488 conjugated rabbit IgG (Jackson ImmunoResearch, West Grove, PA). We also counterstained with 4’,6-Diamidino-2-Phenylindole, Dihydrochloride (DAPI). We digitally scanned the light and fluorescence slides by NDP NanoZoomer XR whole slide scanning system (Hamamatsu Photonics K.K., Japan) and stored as ndpi format for further analysis. We used Imaris Image analysis software (Bitplane USA, Concord, MA) for examining histopathology and influenza A NP stained cells. We exported the analyzed data to Prism 7 software (Graphpad, La Jolla, CA) for statistical analysis.

## Acknowledgements

We thank Adrian McDermott for providing BN HA IgG1 expressing plasmids, Peter Palese for providing cH5/1 expressing plasmids, Kanta Subbarao for providing pHH21-N2 IAV reverse genetic plasmid. We thank Glenys Reynoso for outstanding technical assistance. This work was supported by the Division of Intramural Research of the National Institute of Allergy and Infectious Diseases.

## Author Contributions

Conceptualization, I.K., D.A., M.A., J.W.Y; Methodology I.K., D.A., M.K., H.D.H., J.W.Y.; Software, M.A.; Validation, I.K.; Formal Analysis, I.K., D.A., M.A., K.T., M.K., V.N., J.W.Y.; Visualization, I.K., K.T., M.K., V.N., J.W.Y.; Project Administration, I.K., J.W.Y.; Supervision, J.W.Y.; Funding Acquisition, J.W.Y. H.X., J.W.Y.; Investigation, I.K., J.S.G., D.A., M.A., K.T., M.K., V.N., C.C.B.; Resources, H.D.H., H.X., J.W.Y.; Data curation, I.K., M.A., K.T., M.K.; Writing-Original draft, I.K., D.A.,

---

**Figure Supplementary 1.**
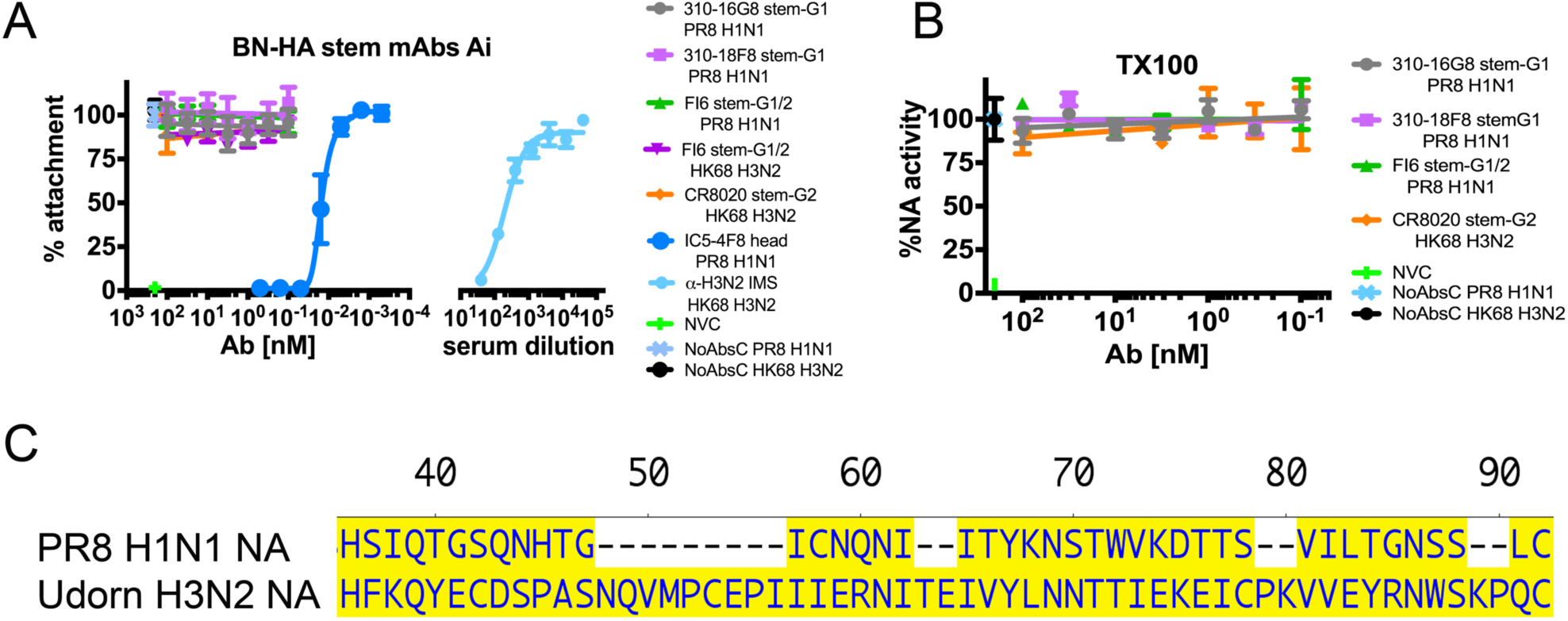
Characterizing anti-stem Ab inhibition of attachment and NA activity. A. We pre-incubated the indicated viruses with the Abs indicated and added to fetuin coated plate at 4°C to enable virus attachment. After removal of free virus we added MUNANA (a fluorogenic substrate for NA) and measured fluorescent signal. Note that stem-Abs do not block attachment, which is efficiently blocked by head specific Abs (IC5-4F8 for PR8, polyclonal mouse serum for HK68). Data are normalized to the fluorescent signal in absence of Ab (set to 100%). Error bars indicate the SD of duplicate samples.
B. We tested BN HA Abs mediated NAI on the HA/NA detergent dissociated virus. TX100 dissociated PR8 H1N1 and HK68 H3N2 viruses were incubated with BN HA Abs. We transferred the mixtures to fetuin coated plate and performed ELLA. We normalized the data by NA activity in the absence of the Ab (set to 100%). Error bars indicate the SD of duplicate samples acquired in two independent experiments. Similar results were obtained in two experiments.
C. We aligned PR8 H1N1 and Udorn H3N2 amino acid sequence of NA stalk region using Lasergene 14/MegAlign software.

**Figure Supplementary 2.**
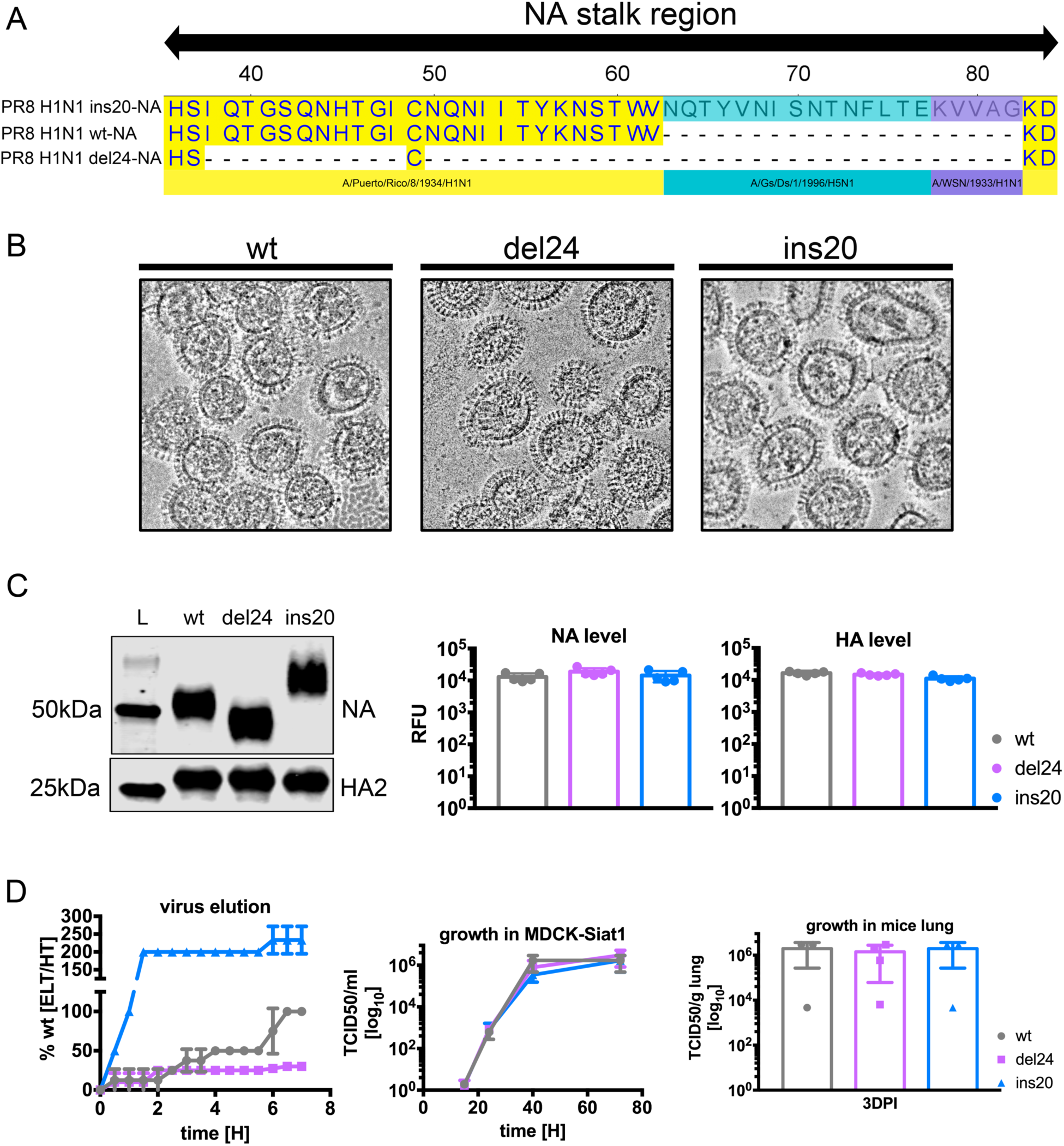
PR8 NA stalk length variants characterization. A. We aligned PR8 H1N1 constructed wt, del24, ins20 amino acid sequence of NA stalk region using Lasergene 14/MegAlign software
B. We preformed cryoEM to compare overall particle shape and size between NA stalk variants.
C. We tested mobility shift of the wt, del24 and ins20 NA in WB and we quantified the signal for NA and HA2 to determine relative amounts of glycoproteins in the virus. Two independent virus preparations were tested in duplicate-triplicates.
D. We compared elution properties of NA variant viruses on RBCs. We normalized viruses by equal NA activity (determined by MUNANA) and maximal elution of wt at 7h was set as 100% elution. Error bars indicate the SD of duplicate samples acquired in two independent experiments. We compared viral growth fitness of NA variant viruses on MDCK SIAT1 cell line (growth kinetics) or in mice (endpoint replication). Error bars indicate the SD of thriplicate-tetraplicate samples.

**Figure Supplementary 3.**
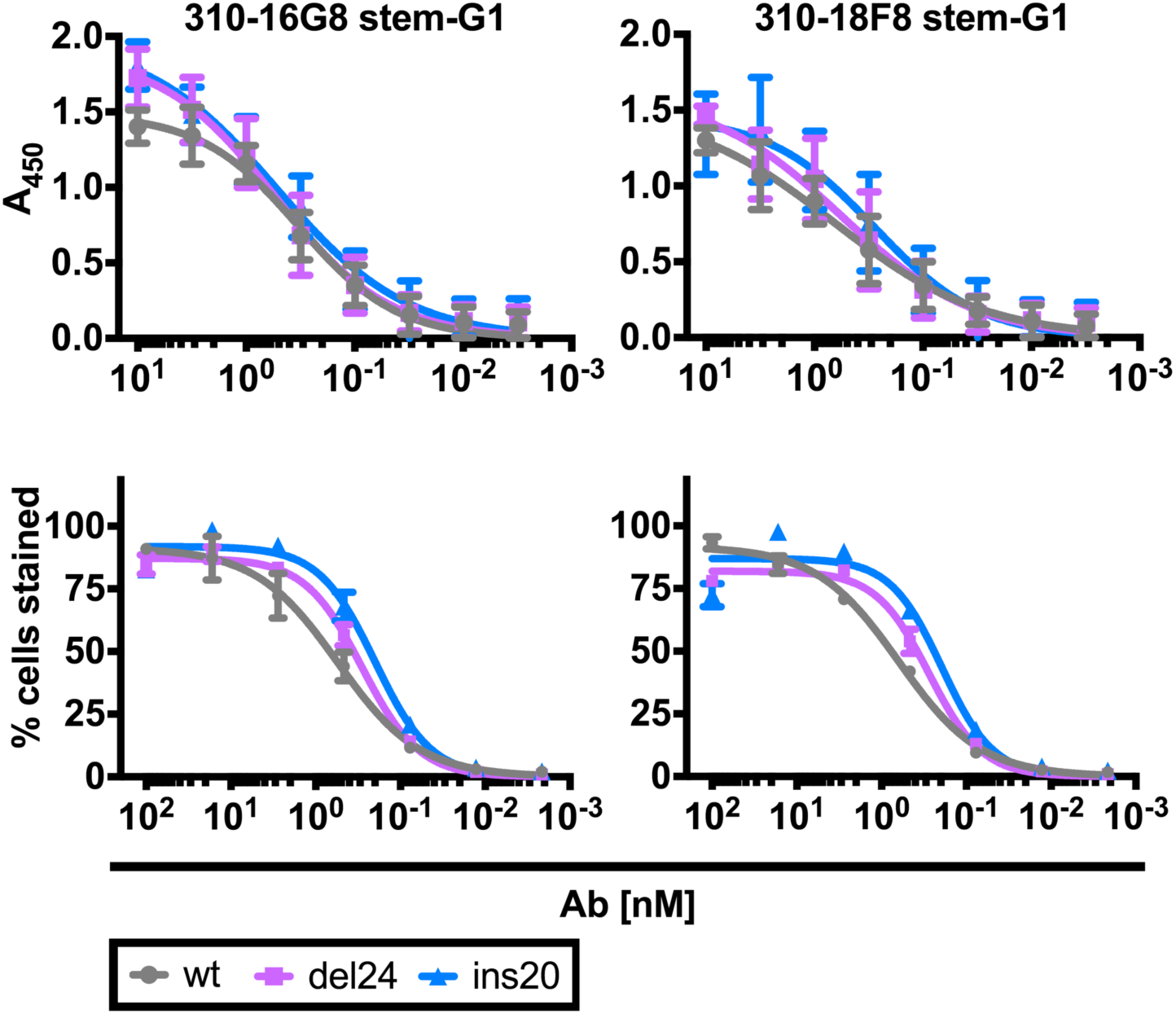
BN HA Abs epitope accessibility. We performed ELISA on amount normalized purified NA variant viruses. We infected MDCK SIAT1 cells with NA variant viruses (moi=1) and at 18 h p.i. we pre-incubated cells with serial dilutions of BN HA Abs. After staining with secondary Ab we measured the positive fraction at given Ab concentration. Error bars indicate the SD of duplicate samples acquired in two independent experiments.

**Figure Supplementary 4.**
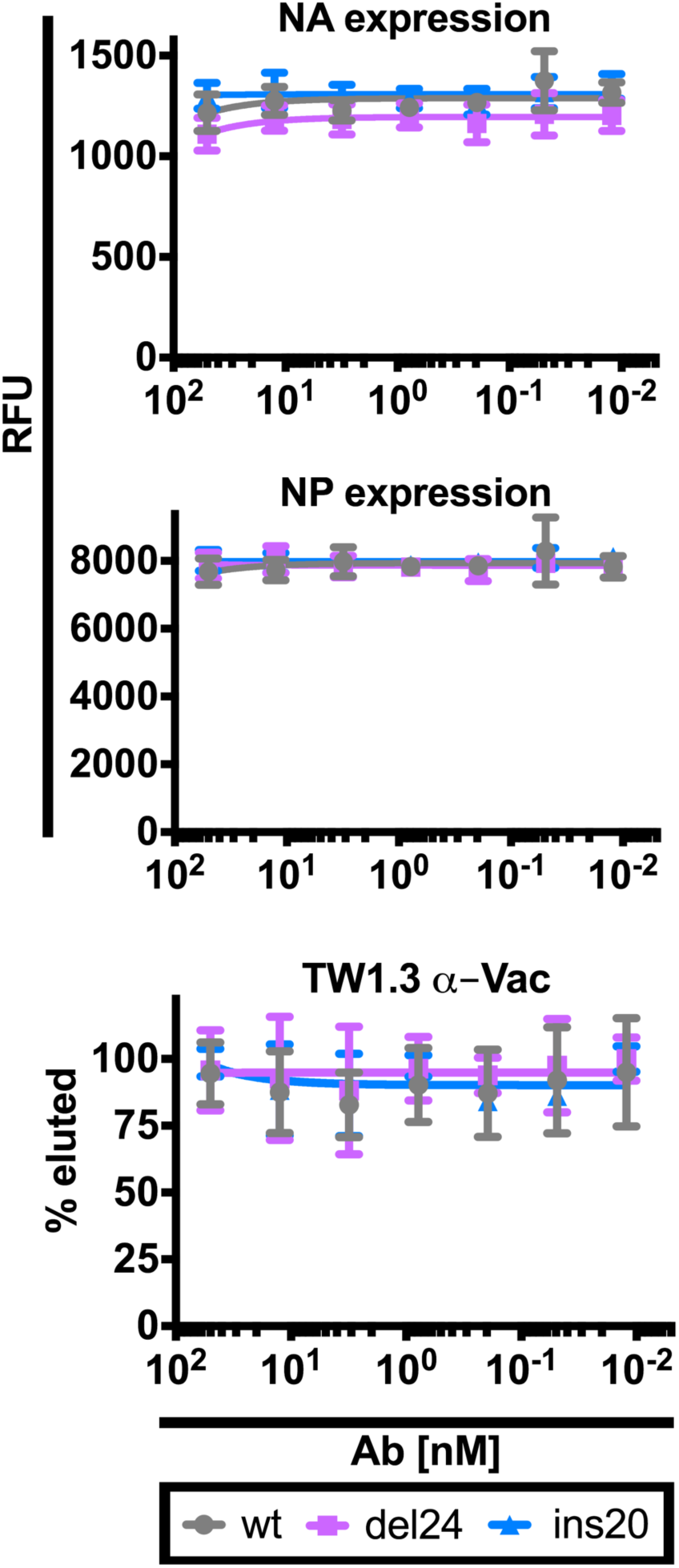
BN HA Abs influence on intracellular and cell surface expression of the viral proteins. We relatively quantified NA and NP expression by indirect immunostaining (in cell western) in MDCK SIAT1 cells treated by BN HA mAbs 4 h p.i. with NA variant viruses. We present combined data for treatment with 310-16G8 and 310-18F8. Error bars indicate the SD of duplicate samples. Similar results were obtained in three to four independent experiments. Four hr p.i. of MDCK SIAT1 cells with the viruses indicated we added graded amounts of the TW1.3 mAb (irrelevant antibody). Six hr later, we collected the supernatant and used NA activity against a fluorescent substrate to quantitate released virions. The fluorescent signal in absence of Ab was set as 100% for each virus. Error bars indicate the SD of duplicate samples acquired in up to four independent experiments.

**Table Supplementary 1.** The NAs enzymatic properties.

| **Michaelis–Menten kinetics** | **wt** | **del24** | **ins20** |
| --- | --- | --- | --- |
| $V_{\max}$ (4-MU [pmol.s<sup>-1</sup>]) ± s.e.<sup>a</sup> | 527.6±8.076 | 406.8±7.652 | 429.2±6.944 |
| Mean $K_m$ (MUNANA [μM]) ± s.e.<sup>b</sup> | 67.09±2.595 | 35.09±2.01 | 60.31±2.545 |
| $V_{\max}$ ratio<sup>c</sup> | 1 | 0.77 | 0.81 |
| $V_{\max}/K_m$ ($E_{\text{eff}}$)<sup>d</sup> | 7.864 | 11.59 | 7.116 |
| $R^2$ goodness of fit | 0.9949 | 0.984 | 0.9936 |

